# Telomere length dynamics in response to DNA damage in malaria parasites

**DOI:** 10.1101/2020.11.08.373563

**Authors:** Jake Reed, Laura A Kirkman, Bjӧrn FC Kafsack, Christopher Mason, Kirk W Deitsch

## Abstract

Malaria remains a major cause of morbidity and mortality within the developing world. Recent work has implicated chromosome end stability and the repair of DNA breaks through telomere healing as potent drivers of variant antigen diversification, thus associating basic mechanisms for maintaining genome integrity with aspects of host-parasite interactions. Here we applied long-read sequencing technology to precisely examine the dynamics of telomere addition and chromosome end stabilization in response to double strand breaks within subtelomeric regions. We observed that the process of telomere healing induces the initial synthesis of telomere repeats well in excess of the minimal number required for end stability. However once stabilized, these newly created telomeres appear to function normally, eventually returning to a length nearing that of intact chromosome ends. These results parallel recent observations in humans, suggesting an evolutionarily conserved mechanism for chromosome end repair.

## INTRODUCTION

Globally, malaria continues to represent a major threat to public health, infecting over 200 million people a year and causing roughly half a million deaths, the majority being children under the age of 5 [1]. The disease is caused by infection with one of several species of eukaryotic parasites of the genus Plasmodium, of which *Plasmodium falciparum* is responsible for the most severe forms of the disease. These parasites infect the circulating red blood cells of their mammalian hosts, resulting in severe anemia, inflammatory problems and disrupted circulation [2]. The complete genome sequence of *P. falciparum* has now been assembled for numerous geographical isolates, providing information into aspects of virulence, mechanisms of drug resistance and the plasticity of the parasite’s genome [3, 4]. In addition, malaria parasites are quite distant from the highly studied model eukaryotes, providing insights into the evolution of many basic molecular processes.

*P. falciparum* has a well-defined chromosome structure that differs somewhat from model eukaryotes. The genome includes 14 chromosomes, and each can be sub-divided into two structural components, the core chromosome containing the centromere and housekeeping genes, and the subtelomeric regions (30-120kb) containing telomere-associated repetitive elements (TAREs), multi-copy gene families encoding variant surface antigens (including *var*, *rifin*, *stevor* and *Pfmc-2TM*) and telomere repeats [5]. These elements are found in a relatively uniform arrangement, with the multi-copy gene families positioned at the boundary of the core genome followed by TAREs and terminating with telomeres at the chromosome ends [3]. This arrangement functionally partitions the genome into the core, which displays near complete synteny and a very high degree of sequence conservation when comparing different geographical isolates, and the hyper-variable, multi-copy gene families that are subject to rapid diversification through frequent ectopic recombination events [3, 6–8]. The TAREs are composed of repeat elements which range in size from 21bp to 164bp [9]. Their function has not been defined, although they are known to transcribe long noncoding RNAs [10, 11] and have been hypothesized to play a role in the subnuclear positioning of the chromosome ends, thus possibly contributing to subtelomeric recombination events [5].

The hyper-recombinogenic properties of the subtelomeric regions containing the multi-copy gene families is key to both the parasite’s survival as well as the virulence of malaria caused by *P. falciparum*. Of the various gene families found within these regions, *var* is the best studied. This gene family is highly dynamic, varying in number from 45 to 90 genes within the haploid genomes of different parasite isolates [3]. Each *var* gene encodes a different form of the surface antigen *Plasmodium falciparum* erythrocyte protein 1 (PfEMP1) which is expressed on the surface of the infected red blood cell and is responsible for parasite sequestration away from the spleen via adhesion to the vascular endothelium. Sequestration within blood vessels is not only one of the main mechanisms for immune evasion, but it is also directly implicated in a number of malaria disease pathologies including cerebral malaria and placental malaria. Importantly, the position of PfEMP1 on the infected cell surface exposes it to the humoral immune system of its host, and infected individuals readily make antibodies that efficiently recognize and destroy infected cells [12]. Thus, to avoid clearance by the antibody response, parasites alternate which *var* gene is expressed, effectively cycling through their repertoire of genes in a process that is regulated epigenetically [13]. This process is referred to as antigenic variation and is dependent on extensive variability between *var* genes to avoid cross-reactive antibody responses to different forms of PfEMP1. Moreover, to avoid pre-existing immunity from previous infections, different parasites circulating within a geographical area must differ substantially from each other, thus providing a strong selection pressure for continuous and rapid *var* gene diversification.

To explore the extent of *var* gene diversity globally, Otto et al. analyzed 714 clinical malaria isolates across 12 countries [3, 4]. They found that the isolates in each country contained between 6-21 shared *var* genes based on sequence homology, indicating that the vast majority (79-94%) of *var* genes in each region are unique [4]. Previous geographical surveys of *var* gene sequences detected a similar degree of diversity [14, 15], indicating that the hyper-recombinogenic properties of the subtelomeric regions are a universal property of *P. falciparum* parasites. The structure of the subtelomeric regions, their positioning within clusters located at the nuclear periphery, and the molecular processes of telomere healing, homologous recombination, and telomere maintenance have all been implicated in the multi-copy gene family diversification process. For example, Zhang et al. recently reported that a single double-strand break (DSB) within a sub-telomeric region can lead to a cascade of recombination between *var* genes on different chromosomes, leading to the creation new chimeric *var* genes through a combination of telomere healing and homologous recombination [16]. This process was also noted in three other clones in previously published data [16, 17], suggesting that this is a common mechanism leading to the diversification of this highly variant gene family. Thus, the rapid generation of antigen diversity appears to be inherently linked to the maintenance of chromosome ends.

The primary function of telomeres is to prevent degradation of genetic material during replication, often described as the end-replication problem [18]. The telomeres of eukaryotic organisms consist of tandem arrays of short repeat sequences incorporated at the chromosome ends by telomerase. This enzymatic activity enables replicating cells to counter the shortening of telomeres during DNA replication, thereby maintaining the stability and integrity of the chromosome ends. In addition, in the event of a chromosome break within a subtelomeric region, telomerase can stabilize the broken chromosome end through *de novo* telomere addition (also called telomere healing), a process recently linked to accelerated variant antigen diversification in *P. falciparum* [16]. Given that destabilization of chromosome ends can lead to recombinational cascades that result in rapid diversification of variant antigen genes [6, 8], and that telomere healing appears to play a key role in this process [16, 19], the unusual nature of Plasmodium telomere maintenance has acquired renewed attention [20]. Such studies could provide insights into how malaria parasites balance the need to maintain genome integrity while also undergoing rapid genetic diversification. Here we used long-read sequencing technology to investigate telomere dynamics in *P. falciparum* and to determine how the DNA damage response influences these dynamics.

## RESULTS

### SMRT sequencing allows for accurate length determination of *Plasmodium falciparum* telomeres

*P. falciparum* telomeres are highly variable in length, ranging in size from 0.5-6.5 kb [21] and are composed of a 7bp repeat which varies between the sequence 5’-TTCAGGG-3’ and 5’-TTTAGGG-3’ [22]. Given that telomeres are continuously turning over as part of chromosome replication, their length can vary depending on telomerase activity and recruitment of the enzyme to specific chromosome ends. The consequences of differing telomere lengths are not well understood, although recent work has shown that environmental conditions can influence telomere length distribution [23]. In the past, telomere lengths have been estimated by pulse field gel electrophoresis [24–26], quantitative real-time polymerase chain reaction [27, 28] or in situ hybridization [29, 30], with each method providing an estimate of the number of repeats found at the chromosome ends. However, new long-read sequencing methods, called SMRT (single molecule real-time) sequencing [31], are capable of extending through complete arrays of telomere repeats, thus potentially providing a new method to directly measure telomere lengths at each individual chromosome end, and to detect changes in telomere length caused by alterations in environmental conditions. We explored the utility of this method by applying it to the study telomere dynamics in *P. falciparum*.

In a recent study of DNA repair in response to DNA damage, we applied SMRT sequencing technology to generate *de novo* sequence assemblies from cultured parasites [19], however the focus of that analysis was on structural changes and telomere length was not examined. In the current study, we resequenced the same parasite lines in order to better resolve the genome and investigate telomere length dynamics. Two independent parasite lines, one grown under normal culture conditions and the other exposed to DNA damage through irradiation, were studied. Parasite DNA was isolated using phenol-chloroform extraction [32] and sequenced using SMRT technology from Pacific Biosciences (PacBio). PacBio was chosen as the sequencing platform because it produces reads in the 10s of kilobase range as opposed to 50-250 base pair range as is typical for most next generation sequencing platforms. This allows for greater resolution of telomere tracts. A custom Bash [33] script was then used to retrieve all reads that contained the *P. falciparum* telomere repeat motif (5’-TTTAGGG-3’ and 5’-TTTTGGG-3’) which was followed by calculating the percent composition of telomere repeats in 200bp windows within each read. Quality control, length assessment and chromosome end assignment were performed using a custom R [34] script and minimap2 [35]. Of the 31220 reads containing the canonical *P. falciparum* telomere repeat motifs, we were able to assign 24919 (79.8%) reads to their respective chromosome end. Of the 56 predicted telomere tracts from the two samples, we were able to confidently reconstruct 54 unique telomere tracts with a mean coverage depth of 460 ± 168 (Figure 1A). As expected, the 3’ ends of chromosomes 12 and 13 were absent in the irradiated parasites. This is due to a previously documented event where the 5’ end of chromosome 9 was duplicated onto the 3’ end of chromosome 12 and 13 (Figure 1A). The evidence for this event is discussed in detail in a previous publication [19].

**Figure 1:**
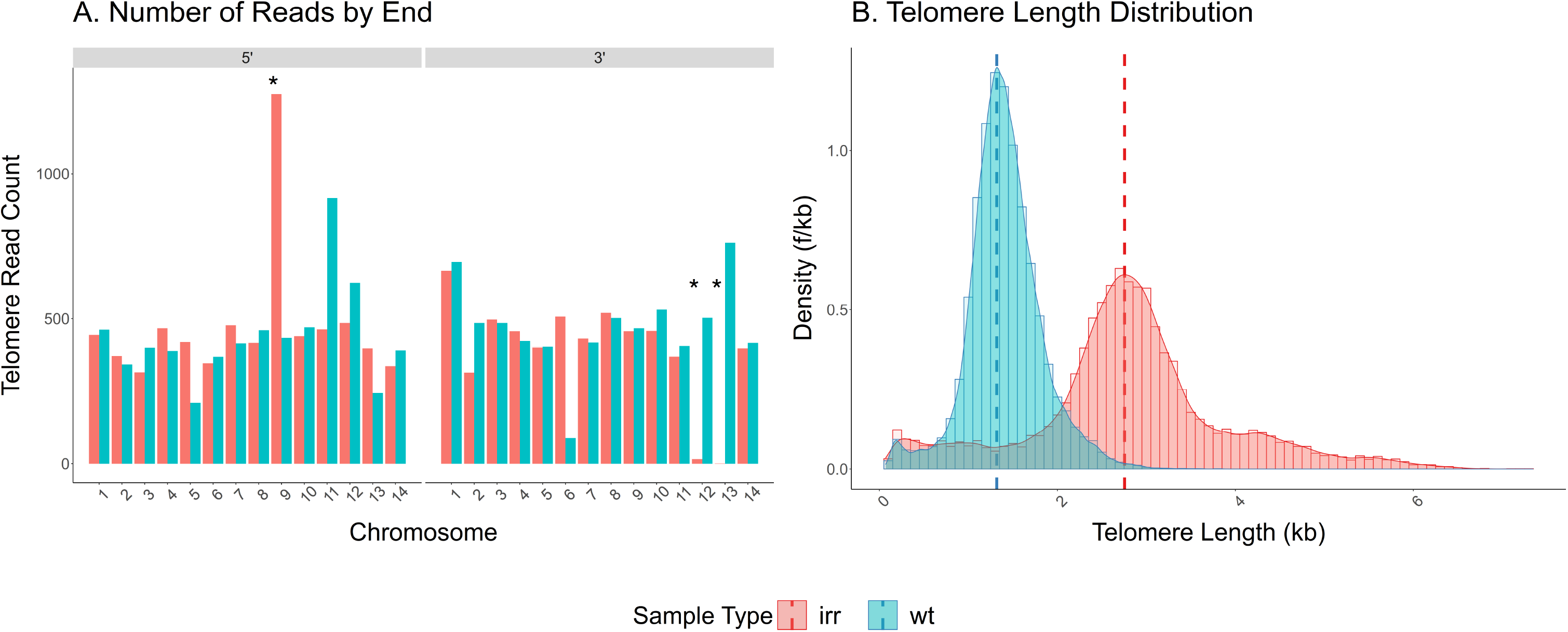
Total telomere length distributions in irradiated and wild-type parasites. (A) Number of telomere reads assigned to each chromosome end by irradiation status, irradiated (red) or wild-type (blue). (B) Comparison of telomere length distributions of the irradiated (red) and wild-type (blue) samples. The red and blue dashed lines indicate the modes of the irradiated and wild-type samples, respectively. (*) Designates a duplication of the 5’ end of chromosome 9 onto the 3’ ends of chromosomes 12 and 13 [19].

In order to determine telomere length, a custom R [34] script was used to estimate length based on a 200bp window falling below a telomere motif content threshold of 60% at which point the number of 200bp windows were counted. This allowed us to resolve telomere length to within 100bp on a per read basis. The telomere length distribution was then plotted and the modes were determined using the modes R package [36]. Thus, our ability to use long-read sequencing technology combined with a computational approach which measured telomere length at 100bp resolution per read allowed us to confidently determine telomere length distributions in *P. falciparum*.

### Irradiated malaria parasites have longer telomeres

In our previous study, we examined DNA repair within subtelomeric regions of *P. falciparum* using X-ray irradiation as a source of DNA damage [19]. This study identified both telomere healing and homologous recombination as important mechanisms for maintaining genome integrity in response to DNA double strand breaks, however how these repair events influenced telomere length was not examined. Given that the repair events that we previously described frequently involved telomere healing, it was clear that telomerase activity was likely influencing the repair process. This led us to investigate if such mechanisms of repair might alter the telomere lengths observed in these parasites. We therefore more precisely examined dynamics specifically at the chromosome ends by directly comparing telomere length distributions in genomic DNA extracted from both the irradiated and non-irradiated parasite cultures.

For the irradiated samples, mixed stage parasites were exposed to X-rays at 100 Gy for three iterations, allowing the parasites to regrow in between each exposure [19]. After recovery, DNA was isolated from a clonal line of parasites obtained from the irradiated population, sequenced using PacBio technology and analysed as described above. When the telomere lengths derived from both the irradiated and non-irradiated samples are displayed side by side, the telomeres from the irradiated sample displayed a clear increase in average length (p <0.001) (Figure 1B). The mean length for the wild-type parasites was 1.409 kb while the mean length of the irradiated parasites was 2.806 kb, which amounts to a near doubling of general telomere length due to exposure to x-rays (Figure 1B).

### Sites of telomere healing have significantly increased numbers of telomere repeats

We were curious if the increased average telomere length observed in the irradiated line was due to a general increase in the number of telomere repeats across all chromosome ends, or if instead specific telomeres displayed disproportionate increases in length. We were especially interested in sites of recent telomere healing where increased recruitment and processivity of telomerase are required for synthesizing a new telomere *de novo*. The unique structure of *P. falciparum* subtelomeric domains makes identifying healing events simple and unambiguous. All wildtype subtelomeric domains include the telomere repeats at the chromosome end flanked by 10-25 kb of TAREs followed by members of the multi-copy variant antigen gene families. The loss of TAREs and/or insertion of telomere repeats within a region containing multi-copy gene family members is a hallmark of telomere healing, with the site of telomere addition easily identifiable as the position where the telomere repeat sequences initiate and the chromosome prematurely terminates. Further, by comparing genome assemblies before and after exposure to radiation, we can identify telomere healing events that were present in the parasite’s genome prior to exposure to irradiation (past events) and recent events resulting from radiation exposure.

Our previous analysis identified six sites of telomere healing that existed in our parasite line prior to radiation exposure and two new sites of healing that were the direct result of exposure to irradiation, the 5’ end of chromosome 1 and the 3’ end of chromosome 2 [19]. Our current analysis also identified a third radiation induced telomere healing event on the 5’ end of chromosome 3 (Figure S1). Further, since the DNA for analysis was isolated from parasites shortly after they had recovered from radiation exposure, we had the unique opportunity to measure telomere lengths at sites of relatively recent telomere healing events. Thus, by comparing the telomere lengths of all 28 chromosome ends for both the irradiated and non-irradiated lines, we could examine telomere dynamics as parasites recovered from DNA damage.

Individual comparisons of the telomere lengths of each chromosome end detected a general trend toward longer telomeres at all chromosome ends in the irradiated parasite line (Figure 2), with recently healed telomeres displaying particularly pronounced increases in length (Figure 2, red boxes). This suggests that the DNA damage induced by exposure to irradiation led to a general increase in telomerase activity, thus leading to an increase in telomere lengths at all chromosome ends. However, the recruitment of telomerase to sites of telomere healing induces the initial synthesis of telomere repeats well in excess of the minimal number required for end stability, thus leading to the particularly long tracts of telomere repeats observed at healed chromosome ends. This increased length was sufficiently stable that it was easily observed after the several months in culture required for recovery from irradiation and for expansion of the parasite population after cloning.

**Figure 2:**
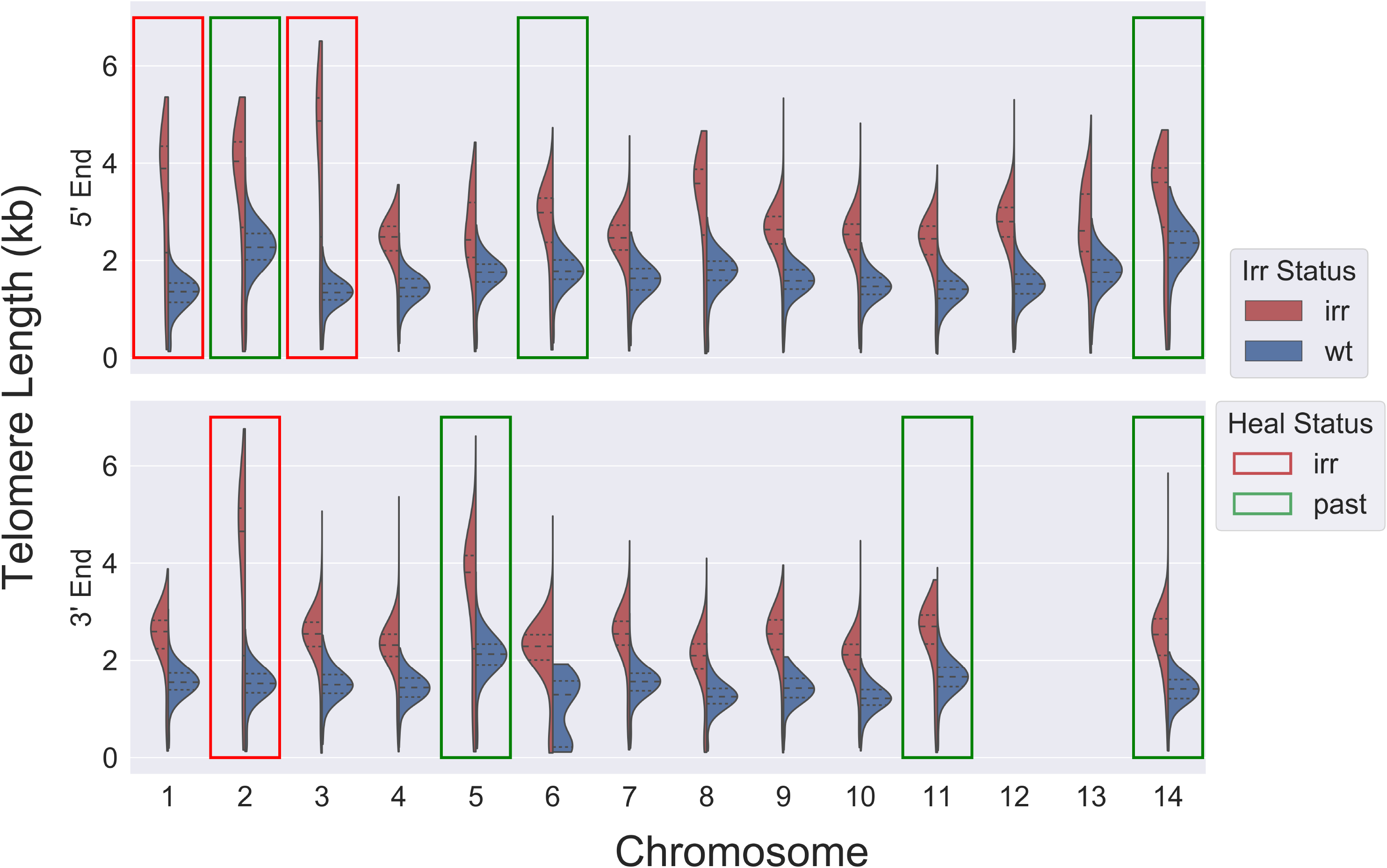
Comparison of telomere lengths by end and irradiation status. Red boxes indicate telomere healing events resulting from exposure to irradiation while green boxes indicate telomere healing events found in the original population, prior to exposure to irradiation.

### Lengthened telomeres at sites of healing shorten over time, but remain extended

Our examination of the recently irradiated parasite line indicated that the changes in telomere repeat numbers were relatively stable, at least over the course of the several months required to complete the experiment. This raised the possibility that the increased telomere length at sites of healing might be required to compensate for the altered structure of the subtelomeric domain, for example due to the loss of TAREs after healing, and therefore permanent. To investigate this possibility, we took advantage of our recent identification of six sites of telomere healing that were present in the original population of parasites prior to irradiation (Figure 2, green boxes) [19]. These healing events occurred sometime in the history of this parasite line, and five are found in the reference sequence of 3D7 (Eupathdb), indicating they likely occurred prior to the isolation of the 3D7 clone over 30 years ago. This enabled us to compare chromosome healing events that happened in the distant past to very recent events and thus infer details of telomere length dynamics and stability in malaria parasites. To facilitate this analysis, we grouped telomere tracts into bins based on healing status and whether the parasites had been exposed to irradiation, then compared the length differences of each class using Welch’s t-test to evaluate the significance of changes (Figure 3).

**Figure 3:**
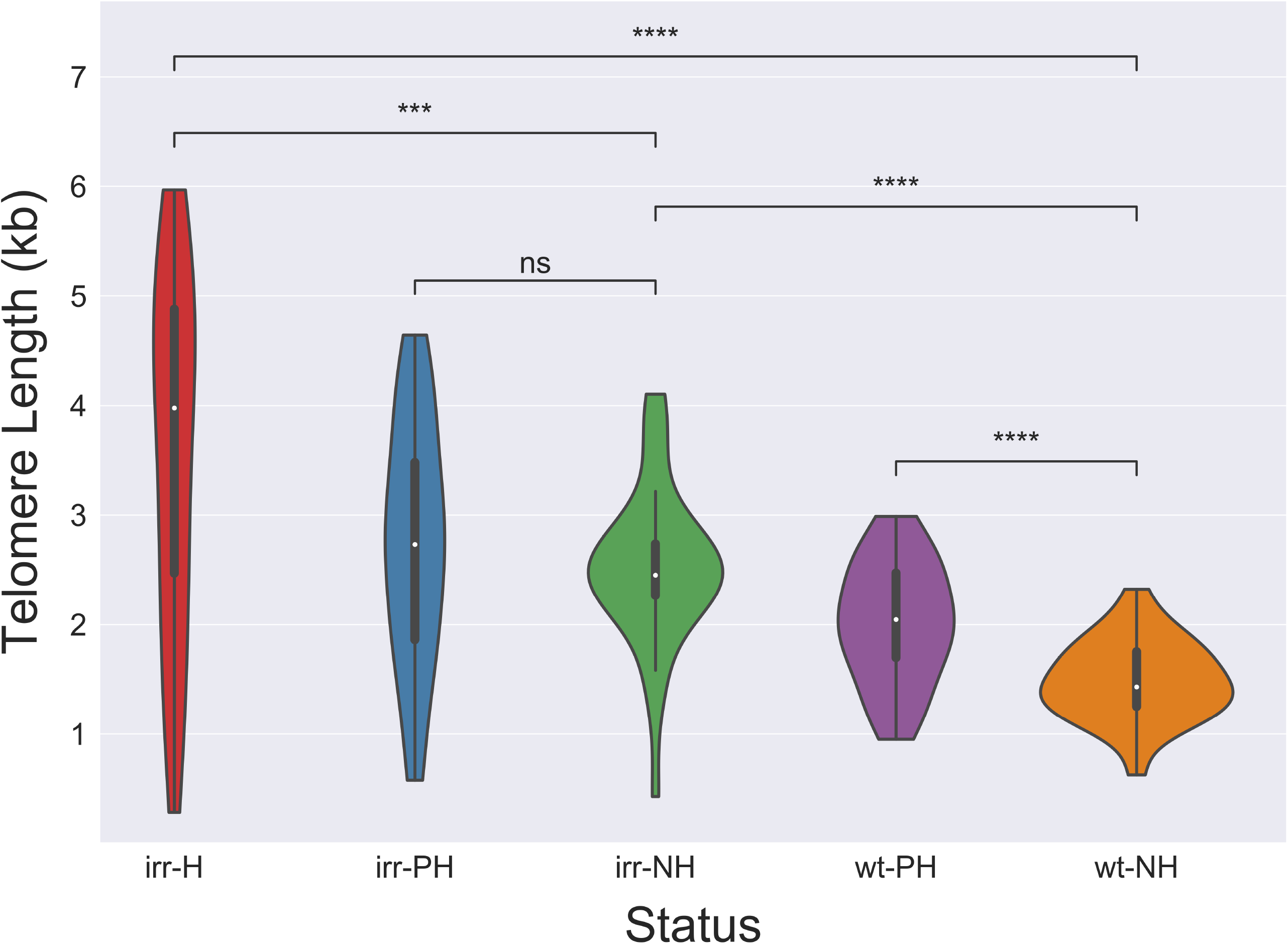
Telomere Lengths by Irradiation and Healing Status. Telomere lengths were assessed after a random downsampling to n=35. Welch’s t-test was used to determine significance. The samples are ordered from left to right, irradiated healed telomere tracts (irr-H), irradiated past healed telomere tracts (irr-PH), irradiated non-healed telomere tracts (irr-NH), wild-type past healed telomere tracts (wt-PH), and wild-type non-healed telomere tracts (wt-NH). p-value annotation legend: ns: 5.00e-02 < p <= 1.00e+00 *: 1.00e-02 < p <= 5.00e-02 **: 1.00e-03 < p <= 1.00e-02 ***: 1.00e-04 < p <= 1.00e-03 ****: p <= 1.00e-04

When telomeres from irradiated parasites were compared to non-irradiated controls, non-healed telomeres (irr-NH) increased in length by 1.033 kb (1.69 fold increase) while the healed ends (irr-H) displayed a much greater increase of 2.072 kb (2.39 fold increase) (Figure 3 and [Table 1]). As described above, this is consistent with a general increase in telomerase activity in response to irradiation leading to a lengthening of all chromosome ends, with an even greater increase in telomerase activity during *de novo* telomere addition. The significance of these events by Welch’s t-test were p < 0.001 after downsampling to 35 reads per bin (Figure 3).

**Table 1:** Telomere length increases. The samples are grouped in the left column by irradiation status (wild-type, wt; irradiated, irr) and healing status (recently healed due to irradiation, H; pre-radiation healing, PH; not healed, NH).

| Status (Rad-Healing) | Percentiles (kb) | | | Mean ± SE (kb) | Fold Increase |
| --- | --- | --- | --- | --- | --- |
|  | 25<sup>th</sup> | 50<sup>th</sup> | 75<sup>th</sup> |  |  |
| *n* = 35 |  |  |  |  |  |
| irr-H | 2.471 | 3.979 | 4.879 | 3.560 ± 0.271 | 2.39 |
| irr-PH | 1.861 | 2.733 | 3.479 | 2.747 ± 0.184 | 1.85 |
| irr-NH | 2.266 | 2.452 | 2.737 | 2.521 ± 0.119 | 1.69 |
| wt-PH | 1.698 | 2.046 | 2.472 | 2.012 ± 0.092 | 1.35 |
| wt-NH | 1.251 | 1.431 | 1.753 | 1.488 ± 0.061 | 0.00 |
| *n* = 1075 |  |  |  |  |  |
| irr-H | 2.196 | 4.129 | 4.965 | 3.571 ± 0.052 | 2.48 |
| irr-PH | 2.394 | 2.968 | 3.799 | 2.906 ± 0.035 | 2.02 |
| irr-NH | 2.115 | 2.449 | 2.800 | 2.362 ± 0.023 | 1.64 |
| wt-PH | 1.541 | 1.923 | 2.272 | 1.907 ± 0.017 | 1.33 |
| wt-NH | 1.244 | 1.441 | 1.656 | 1.439 ± 0.011 | 0.00 |

To investigate what happens to healed telomeres over time, we used data from the non-irradiated parasites to compare telomere repeat lengths at sites of past telomere healing events to non-healed telomeres (wt-PH vs wt-NH). This also revealed a significant, albeit more modest increase in telomere length of 534bp (p < 0.001), suggesting that the increased telomere length due to healing eventually returns to a length approaching that of non-healed chromosome ends, but that they remain distinctly longer. Interestingly, a similar comparison of previously healed and non-healed telomeres from the irradiated parasites (irr-PH vs irr-NH, Figure 3) indicated that these lengths were not significantly different, suggesting the possibility that exposure to irradiation leads to the increase of all telomeres to similar lengths (previously healed or non-healed), but that the healed telomeres remain somewhat longer after recovering to a “normal” set point (Figure 3).

The trends described above are evident across all sample sizes (35-1075) indicating consistent differences in this data set (Figure 4). Across all sample sizes, the most significant length differences are between the irr-H vs wt-NH and irr-NH vs wt-NH comparisons whose p values continue to separate further from the other comparisons as sample number increases, indicating there are consistent and highly significant increases in lengthening of telomeres due to healing and exposure to irradiation (Figure 4).

**Figure 4:**
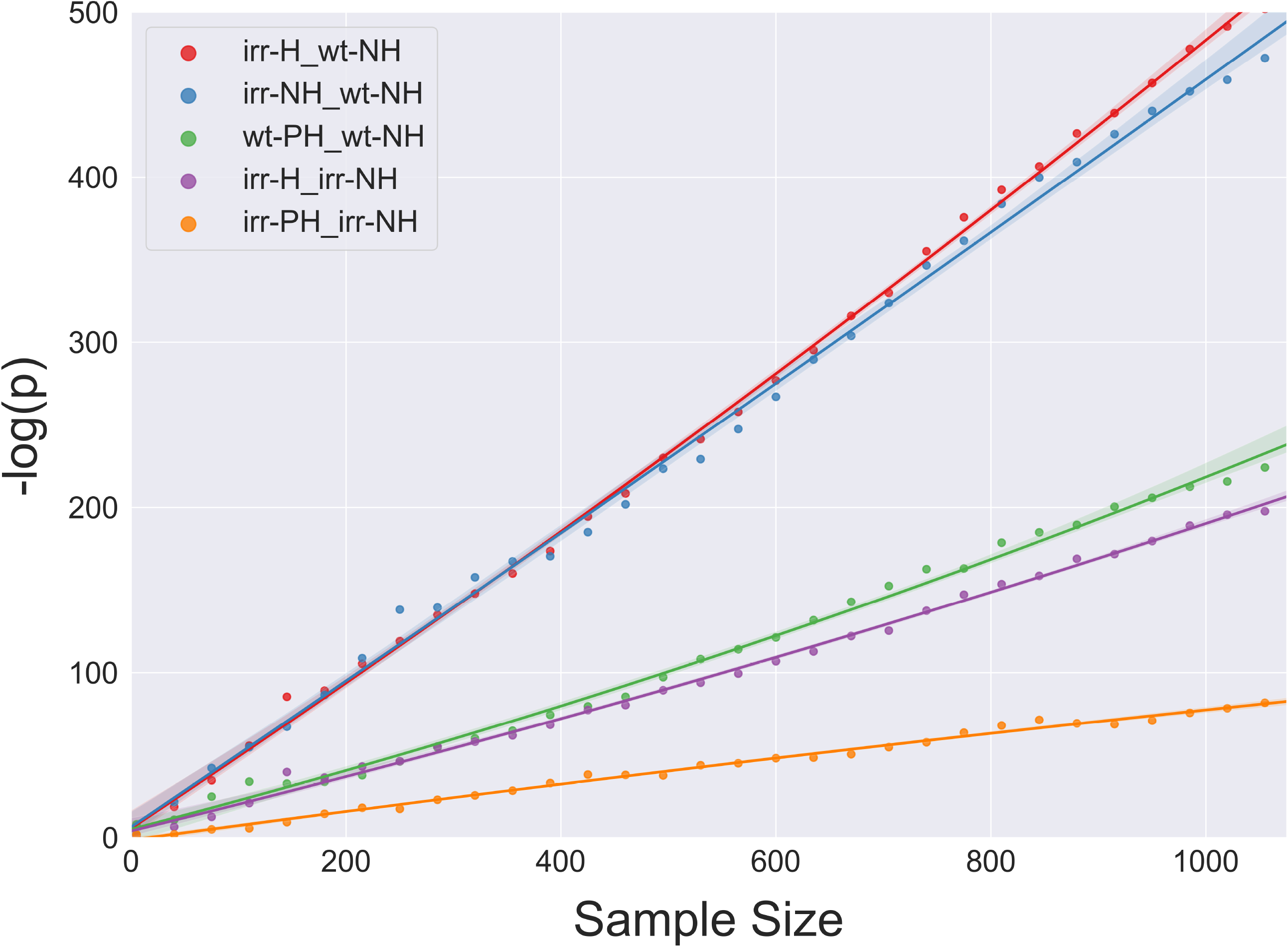
Sample size impact on p-value. Sample size was randomly downsampled from the minimum number of reads per grouping (1075 reads in the irradiated healed set). P-values were generated based on different grouping comparisons while sample size was progressively increased from *n*=35. The p-values generated approached zero in an exponential fashion, therefore −log(p) was plotted in order to make differences discernable. Irradiated healed telomeres vs wild-type non-healed (irr-H_wt-NH) (red) and irradiated non-healed telomeres vs wild-type non-healed telomeres (irr-NH_wt-NH) (blue) comparisons consistently displayed the most significant differences in telomere length. Wild-type past healed telomeres vs wild-type non-healed telomeres (wt-PH_wt-NH) (green) and irradiated healed telomeres vs irradiated non-healed telomeres (irr-H_irr-NH) (purple) displayed marginally significant telomere length differences, and the length difference between irradiated past healed telomeres and irradiated non-healed telomeres (irr-PH_irr-NH) (orange) was non-significant initially but became more significant as sample size increased.

## DISCUSSION

Like in all eukaryotic organisms, telomeres are required for maintaining chromosome stability and genome integrity in malaria parasites. However, in these parasites they also contribute to a unique subtelomeric structure that includes the primary antigenic and virulence determinants of malaria caused by *P. falciparum*. Moreover, telomere healing was recently shown to be a primary contributor to the recombination process that drives the sequence diversification of subtelomeric variant antigen encoding genes [16], thus the synthesis and stability of telomeres directly contributes to host-parasite interactions and the virulence of the disease. Telomere healing has been found in *P. falciparum* field isolates [24, 37, 38], indicating that this mechanism of chromosome stabilization occurs frequently in a natural setting and is not just a phenomenon observed in laboratory cultured parasites. In fact, a common telomere healing event that leads to loss of a gene encoding a protein used in rapid diagnostic tests was recently found to be responsible for decreased efficacy of clinical diagnosis, reinforcing the importance of this process [39, 40]. Therefore understanding the enzymatic activities underlying telomere synthesis and maintenance will provide insights into an important biological process of malaria parasites.

The role of telomere healing in driving *var* gene diversification was recently described by Zhang et al [16]. Their model proposes that when a double strand break occurs within a subtelomeric region, repair of the break by homologous recombination competes with telomere healing to stabilize the chromosome end. When healing occurs, this liberates a hyper-recombinogenic free DNA fragment that often contains *var* genes. This fragment can initiate a cascade of recombination events leading to the generation of new, chimeric *var* genes, thus increasing antigen diversity. Increased telomerase activity, for example as observed here after exposure to radiation, would presumably skew such repair events toward telomere healing and thus accelerate the diversification process. Therefore the basic biology of telomere maintenance and chromosome end repair in *P. falciparum* plays a direct role in pathogenesis and immune evasion.

SMRT sequencing provides a novel method for assaying changes in telomere length with remarkable precision. In addition, as opposed to some other methods, SMRT sequencing also enables easy detection of telomere lengths at individual chromosome ends, thereby allowing the detection of changes that occur only at specific regions of the genome. This enabled us to directly measure the number of repeats resulting from a specific telomere healing event, something that was not previously possible. Using this novel approach, we were able to determine that telomere healing events in *P. falciparum* lead to initial over-lengthening of the telomere, resulting in disproportionately long stretches of telomere repeats at sites of healing. We were also able to determine that the increased length is not permanent and that the extended telomeres appear to eventually return to near baseline. This parallels recent observations in humans in which extended occupation of the international space station led to a temporary increase in telomere length in nucleated blood cells [23]. The telomere lengths returned to baseline after the subject returned to earth. While the cause of the increased telomere length cannot be known for certain, it is tempting to speculate that exposure to increased radiation while in space led to telomere lengthening, similar to what we observed after irradiation of malaria parasites. This suggests a shared response to DNA damage and repair of broken chromosome ends that is conserved across the broad evolutionary distance that separates humans from protozoan parasites.

SMRT sequencing is becoming much more commonly used for the assembly of full genome sequences in a variety of organisms. Given that the sequencing reads included in all of these datasets likely include the telomeres, it should be possible to derive telomere length assessments directly using the methods described here without additional experimentation. Thus, ever increasing application of SMRT sequencing is likely to provide an influx of new datasets with valuable information for researchers interested in the dynamics of telomere maintenance under a variety of conditions. For malaria parasites, our analysis provides additional insights into the unique structure of the chromosome ends and how they are maintained. Given the importance of telomeres and chromosome end stability to variant antigen diversity and expression, the data and methods presented here add to our increasing understanding of this important aspect of Plasmodium biology.

## Acknowledgements

The Department of Microbiology and Immunology at Weill Medical College of Cornell University acknowledges the support of the William Randolph Hearst Foundation. This work was supported by the National Institutes of Health (AI 52390 to KWD, AI 99327 to KWD and LAK, AI76635 to LAK, AI141965 to BFK). KWD is a Stavros S. Niarchos Scholar and a recipient of a William Randolf Hearst Endowed Faculty Fellowship. LAK received support from the William Randolph Hearst Foundation as a Clinical Scholar in Microbiology and Infectious Diseases. We would also like to thank funding from the Bert L and N Kuggie Vallee Foundation to CM, the WorldQuant Foundation to CM, The Pershing Square Sohn Cancer Research Alliance to CM, NASA (NNX14AH50G) to CM, the National Institutes of Health (R01ES021006, 1R21AI129851,1R01MH117406) to CM, TRISH (NNX16AO69A:0107, NNX16AO69A:0061) to CM, The Bill and Melinda Gates Foundation (OPP1151054) to CM, and the Leukemia and Lymphoma Society (LLS) grants (LLS 9238-16, LLS-MCL-982) to CM. The funders had no role in the study design, data collection and analysis, decision to publish, or preparation of the manuscript.

## Author Contributions

JR performed the experiments, collected and analysed the data. BFK aided in designing custom scripts for data analysis. LAK, KWD and CM designed the experiments and analysed data. JR, KWD, LAK and CM wrote the paper.

## METHODS

### Culturing and irradiation

The method for parasite culturing is based on the original method described by Trager and Jensen with modifications as described in Calhoun et al. [41]. Briefly, *P. falciparum* was cultured in RPMI complete media supplemented with Albumax II and gentamicin. The cultures were maintained in a 90% nitrogen, 5% carbon dioxide, 5% oxygen atmosphere at 37*°* Celsius. The ionizing radiation dose was 100-Gy, and mixed-stage parasites were consecutively irradiated three times. The cultures were allowed to recover to normal growth levels between each irradiation exposure. Deletion of *var* genes was assayed by quantitative polymerase chain reaction (PCR) with genomic DNA (gDNA) using a PCR *var* panel developed by Salanti and colleagues [42]. The whole-genome analysis was performed on a subclone chosen due to the loss of three subtelomeric *var* clusters based on the previously described assay.

### Genomic DNA isolation and PacBio Library Preparation

This technique was previously described in Vembar et al [43]. Parasite gDNA was isolated from cultures at 5-8% parasitemia. The DNA extraction method used was phenol-chloroform and ethanol precipitation. Library preparation, size selection and sequencing carried out at Cold Spring Harbor Laboratories. Briefly, SMRTbell template prep kit 1.0 (Pacific Biosciences) was used for the library preparation along with size selection using BluePippin for 20kb fragments which was then sequenced on the PacBio Sequel II platform. Each pool of three samples was barcoded (Sequel_16_barcodes_v3), pooled for equimolar mixing, and then sequenced on a single SMRTcell. The controls (LID50609-Cell3) yielded 3,033,878 polymerase reads with a mean read length of 58,129nt and a subread N50 of 8,491nt, whereas the irradiated samples (LID50610-Cell1) yielded greater total reads, with a total of 4,986,496 reads, a mean read length of 53,656nt, and a subread N500 of 7,638nt. These methods are described in more detail in Calhoun et al [19]. The genomic DNA samples analysed for this study were originally isolated and described by Calhoun et al [19] and resequenced for the current analysis.

### Extracting telomere reads

The computation method utilized was a custom script written in Bash [33]. It involved pattern recognition based on the previously published *P. falciparum* telomere motifs [22]. The raw PacBio subreads were extracted which had any match to the 7bp motif repeated twice, whether that be the original sequence, the complement or the reverse complement. The irradiated sample had a much higher sequencing throughput, therefore, the irradiated telomere reads were randomly down-sampled to match the non-irradiated sample, 15610 reads. The telomere reads from each respective sample were then split into 200bp windows.

### Assessing telomere reads

Once each read containing the canonical repeats were extracted and split, the telomeric content was assessed. This was done with a custom Bash [33] script which, due to the degenerate nature of *P. falciparum* telomeres, used much more promiscuous search patterns as opposed to the previously known telomere motifs. These patterns contained high GC content or high T content. The percent hit of the pattern was calculated across each 200bp sliding window. From there, the telomere sliding window data was imported into multiple custom R [34] scripts which determined the length and the distribution of telomere lengths, and assigned reads to each chromosome end based on the long-read mapping algorithm minimap2 [35]. Figures were then produced using ggplot2 [44] in R [34] and the python packages statannot [45] and seaborn [46]. The statistics for the violin plots were produced using Welch’s independent t-test for normally distributed data.

## SUPPLEMENTAL DATA

**Supplemental Figure 1:**
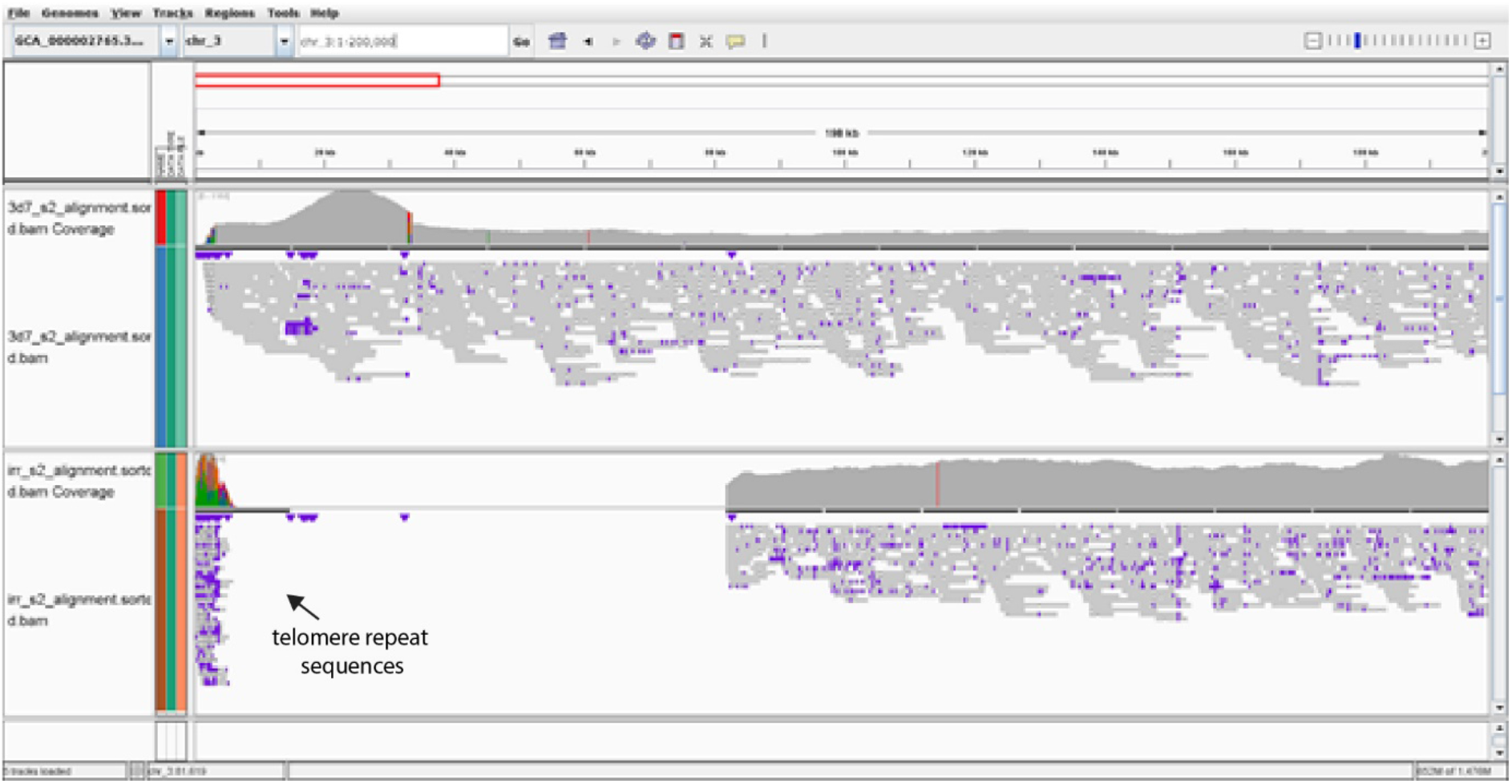
Pile-up of 200kb of the 5’ end of chromosome 3. Circular consensus sequences reads were aligned to the reference genome using minimap 2 [35]. The wild-type parasite reads are on the upper panel while the irradiated parasite reads are on the lower panel. The truncation on the bottom panel (irradiated parasites) “left” end of chromosome 3 is from 1-81,893 bp. The sequence reads from the irradiated parasite include the incorporation of telomere repeat sequences at the site of the deletion, verifying that a telomere healing event has stabilized the chromosome end. This can be observed in the pile-up as reads mapping to the extreme left end of the reference sequence, where telomere repeat sequences reside.

**Supplemental Figure 2:**
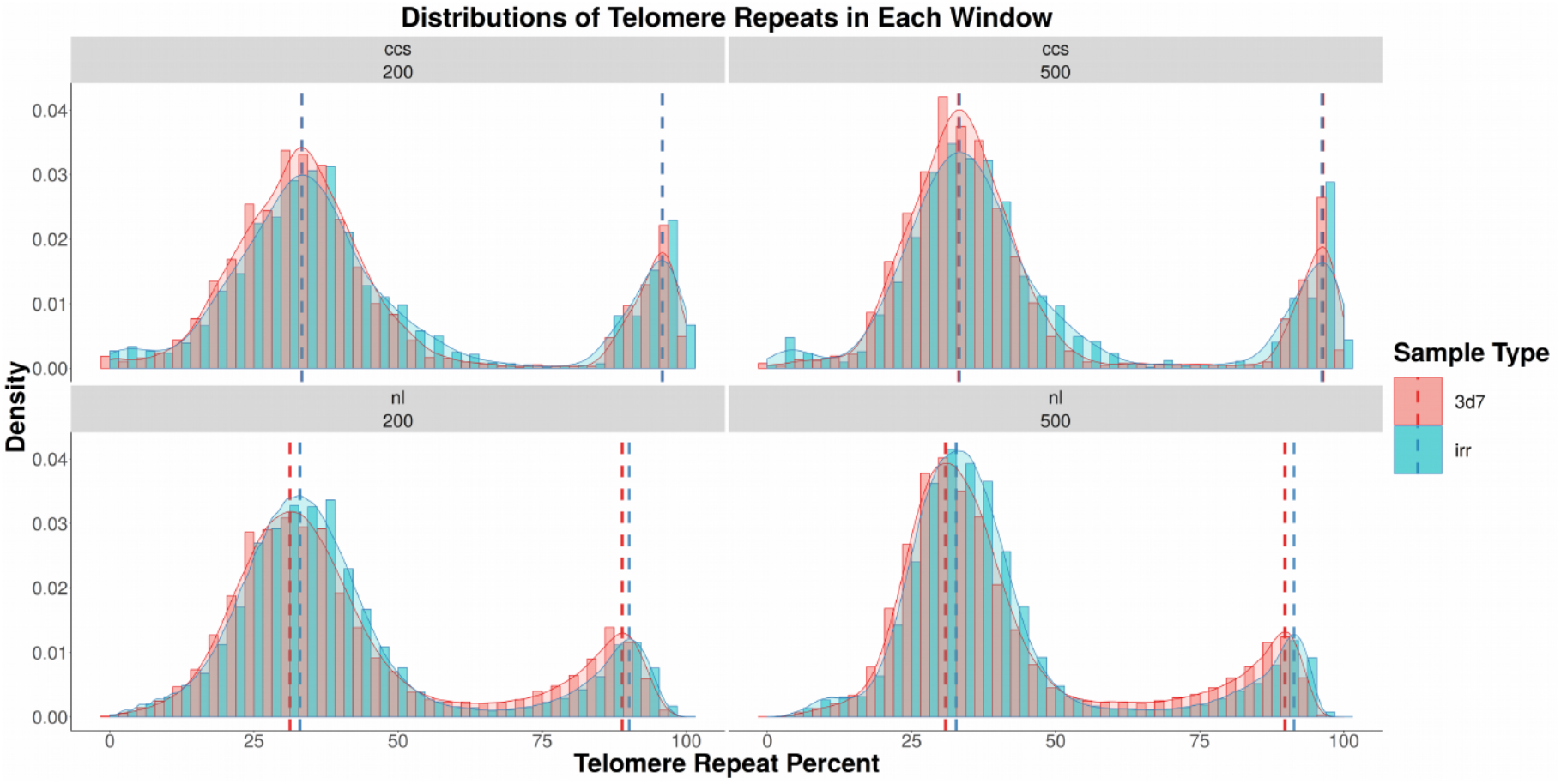
Frequency distributions of telomere repeat percent across each sliding window (200bp, 500bp) and read depth (1x for CCS and 3x for nl). Irradiated samples are in blue (irr) while non-irradiated samples are in red (3d7). The top two figures are circular consensus reads (ccs), and the bottom two figures represent raw pacbio reads or normal (nl). The dotted lines represent the modes in each data set.

**Supplemental Figure 3.**
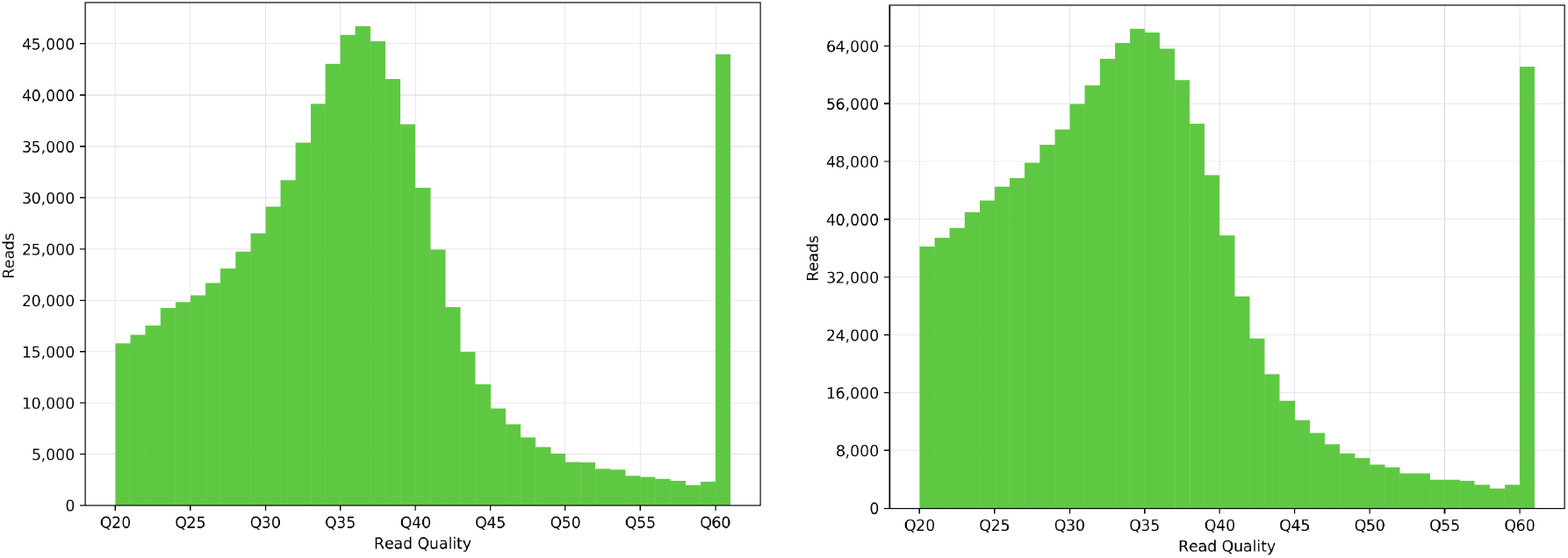
Histogram of Read Quality per Sample Set. The animals for the control samples (left) showed similar distributions of quality metrics as those from the irradiated animals (right). Q-values are shown on the x-axis and the read counts are on the y-axis.

## Notes

### Competing Interest Statement

The authors have declared no competing interest.

